# Reversal of microRNA-blocked tumor suppressors by biphenyldicarboxylate impedes endotoxin-induced hepatic hyperplasia

**DOI:** 10.1101/432856

**Authors:** Li-Li Tan, Dong-Mei Chen, Jian-Ping Song, Qin Xu, Chang-Qing Li, Qing-Ping Zeng

## Abstract

Hepatocellular carcinoma (HCC) is one of the most common malignant tumor and the leading cause of cancer-related death worldwide. Biphenyldicarboxylate (BPDC), an intermediate of schisandrin C from *Schisandra chinensis*, has been used as a hepatoprotective agent that compromises hepatic injuries in China for decades. Whether BPDC is also implicated in the prevention of HCC remains understood. Here, we report that the bacterial endotoxin lipopolysaccharide (LPS) promotes hepatic inflammation and hyperplasia, during which the common tumor markers, alpha fetoprotein (AFP) and carcinoembryonic antigen (CEA), were unregulated, whereas the tumor suppressors, PTEN, FOXO1 and MEN1, were downregulated through increasing the microRNAs, miR-21, miR-122 and miR-24. In contrast, BPDC dampened hepatic inflammation and hyperplasia accompanied by the upregulation of PTEN, FOXO1 and MEN1 through decreasing miR-21, miR-122 and miR-24. However, BPDC failed to downregulate the tumor marker AEG-1 via increasing miR-195. Taken together, BPDC exerts anti-tumor effects by upregulating tumor suppressors upon decreases of miRNAs rather than downregulating tumor markers by increases of miRNAs.

## Introduction

Biphenyldicarboxylate (BPDC or DDB) is an intermediate in the synthesis of schisandrin C in *Schisandra chinensis* (1). BPDC has been widely used for treatment of liver injury with elevated transaminase induced by chemical poisons and drugs from early 70’s in China (2,3). Accumulating data have shown that BPDC exerts a protective role in mice from carbon tetrachloride (CCl4)-induced hepatic injuries, by which the serum aspartate aminotransferase (AST) and alanine transaminase (ALT) are dramatically declined, and hepatocyte steatosis and fibrogenesis are also alleviated (4,5). BPDC was reported to markedly inhibit the inflammatory response by downregulating the expression of TNF-α (6), and to exert a curative effect on tamoxifen-induced liver injuries through scavenging reactive oxygen species (ROS) (7). BPDC was also recognized to enhance liver detoxification and reduce liver damage by activating cytochrome P450 (8). BPDC represses the invasion and metastasis of MHCC97-H cells (10), and downregulates the oncogenic *c-myc* and alpha fetoprotein (AFP), but upregulates the tumor suppressive p53 in Bel-7402 cells (11). However, BPDC seems not to inhibit the replication and expression of HBV in mice (9). These results suggested that BPDC may also play an anti-tumor effect except for amelioration of chemical poison-/drug-induced liver injuries.

It has been reported that many microRNAs (miRNAs) are involved in hepatic diseases, such as autoimmune hepatitis (12), drug-induced liver injury (13,14), alcoholic liver disease (15), non-alcoholic fatty liver disease (NAFLD) (16), liver fibrosis (17,18), and heptocellular carcinoma (HCC) (19,20). The miRNA species are a class of endogenous single-stranded RNA molecules comprising of about 22 nucleotides in length, and can post-transcriptionally regulate gene expression. The altered expression of various miRNAs, including miR-122, miR-195, miR-21 and miR-24, have been documented in HCC development, suggesting an involvement of miRNAs in hepatic pathogenesis (21–24). However, it has not been filed whether BPDC is implicated in the prevention of hepatic hyperplasia by regulating miRNA profiles.

Here, we report for the first time that BPDC impedes hepatic hyperplasia via upregulating tumor suppressors rather than downregulating tumor markers/oncogenes. These results should shed light on exploiting the exact mechanism of BPDC in the prevention and treatment of NAFLD and HCC, and would be also beneficial to elucidating the implication of miRNAs in the development of metastasis of hepatic hyperplasia and other hepatic pathogenesis.

## Materials and methods

### Animals and treatment procedures

Bal b/c mice, belonging to an inbred mouse population, were purchased from the Experimental Animal Centre of Guangzhou University of Chinese Medicine in China (Certificate No. 44005800007512). All mice were housed in a controlled room at the temperature of 24±2°C and the relative humidity of 55%±10% in a 12-h light: 12-h dark cycle with free diet consumption and water drinking. After feeding for one week, mice were randomly divided into three groups: (1) a group of the normal control (NC) of mice; (2) a group of mice administering LPS (LPS); (3) a group of mice given LPS and BPDC (LPS+BPDC). NC mice were intraperitoneally injected with neutral saline. LPS mice were intraperitoneally injected with LPS at 0.25mg/kg every other day for eleven weeks.LPS+BPDC mice were administered BPDC by intragastrical gavage at a dose of 100 mg/kg every day after eight weeks of intraperitoneal injection of LPS. LPS (sigma, L2880) was purchased from Guangzhou HaoMa Biological Technology Company. BPDC, manufactured by Beijing Union Pharmaceutical factory, was purchased from the first affiliated hospital of Guangzhou University of Chinese Medicine. Animal procedures were in accordance with the animal care committee at the Guangzhou University of Chinese Medicine, Guangzhou, China. The protocol was approved by the Animal Care Welfare Committee of Guangzhou University of Chinese Medicine.

### Histopathological analysis

For histopathological evaluations, the liver tissue was fixed in 10% formalin solution and embedded in paraffin. The paraffin-embedded liver tissue was sectioned to 5 µm in thickness, deparaffinized, hydrated, and stained with hematoxylin-eosin (H & E) for histopathological examination under a light microscope.

### Quantitative polymerase chain reaction (qPCR)

The procedure of qPCR includes the following three steps: RNA extraction, reverse transcription, and real-time quantitative PCR. Total RNA was extracted from mouse liver tissue samples by using the trizol reagent. The following thermal cycling conditions for reverse transcription and quantitative PCR are as follows: for RT, 42°C for 60 minutes, 72°C for 15 minutes; for qPCR, 50°C for 2 minutes, 95°C for 2 minutes, and 40 cycles of 95°C for 3 seconds and 60°C for 30 seconds. The U6 was used as a reference gene to analyze the following microRNAs: miR-24, miR-21, miR-195 and miR-122. The sequences of the primers were listed in Table 1. The copy numbers of amplified genes were calculated using the 2^-ΔΔCT^methods.

**Table 1.**
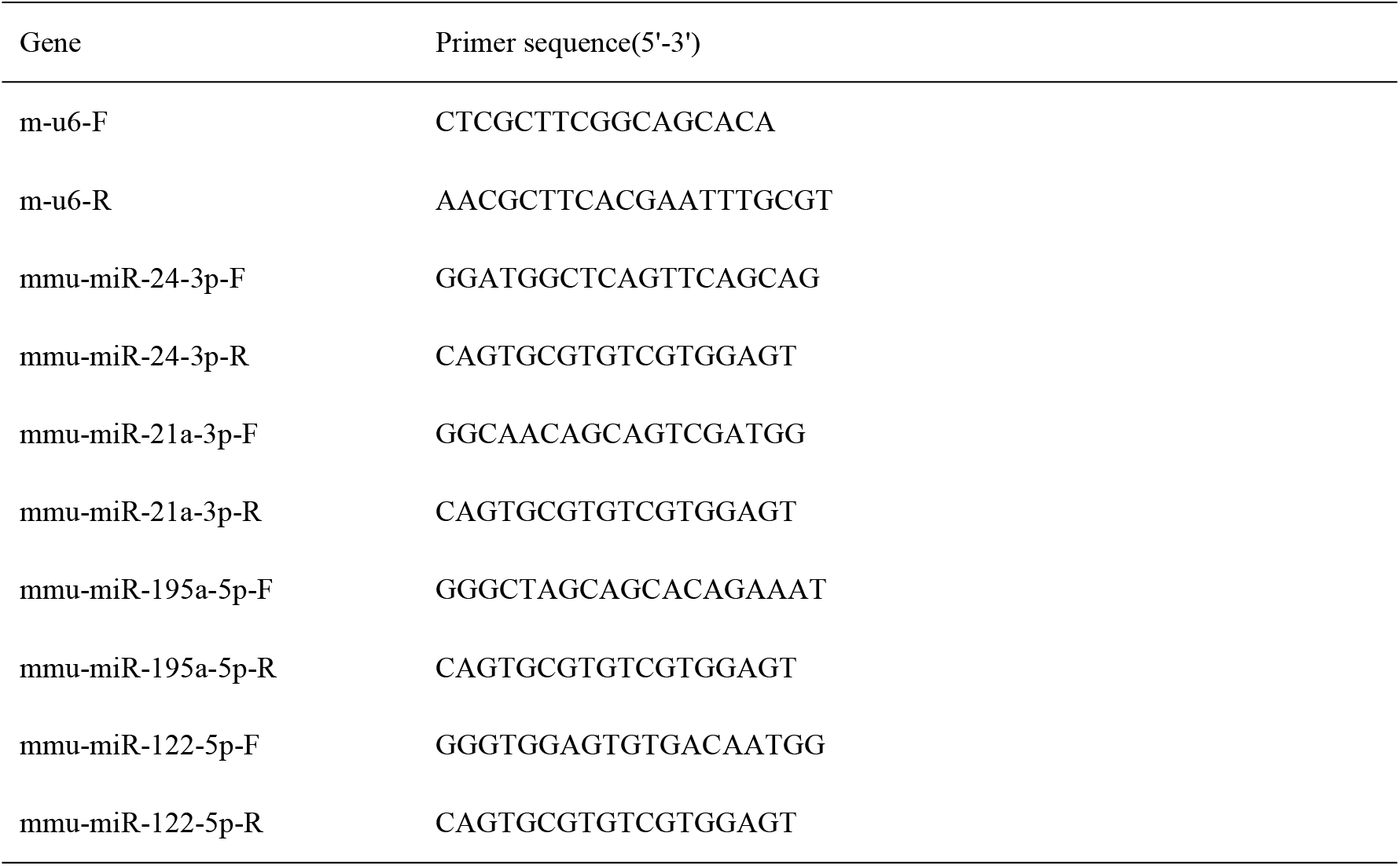
Primer sequences for qPCR.

### Enzyme-linked immunosorbent assay (ELISA)

The ELISA kits were used for determination of pro-inflammatory cytokines, liver cancer-specific tumor markers and corresponding downstream target proteins in the sample of mice. ELISA kits for detecting mouse tumor necrosis factor α (TNF-α), interleukin 6 (IL-6), interleukin 1β (IL-1β), alpha fetoprotein (AFP), carcinoembryonic antigen (CEA), astrocyte elevated gene-1 (AEG-1), multiple endocrine neoplasia 1 (MEN1), phosphatase and tensin homolog deleted from chromosome 10 (PTEN), and Forkhead Box Protein O1 (FOXO1) were purchased from Beijing Chenglin Biotech Company, China. All procedures were operated strictly in accordance with the manufacturer’s manual.

### Statistical analysis

Statistical analyses were conducted by using Graphpad prism 6.01 (GraphPad Software, La Jolla, CA, USA). All date were expressed as 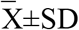 unless otherwise stated. Statistical significance was analyzed by two-tailed Student’s t test between two groups. Experiments with more than three groups were analyzed with one-way analysis of variance (ANOVA) and Bonferroni test. It is generally considered that a *P*<0.05 value is statistically significant, and *P*<0.01 is statistically very significant.

## Results

### LPS upregulates liver cancer-specific tumor markers

It is well known that LPS activates the pattern recognition receptor, toll-like receptor 4 (TLR4), and conveys the inflammatory signal to macrophages, mainly kupffer cells in the liver, causing the release of pro-inflammatory cytokines, such as IL-6, IL-1β and TNF-α. We tried to assess whether LPS-induced TNF-α, IL-1β and IL-6 would be correlated with expression of the specific liver cancer markers, AFP and CEA in mice. For this purpose, we exposed mice to LPS and then measured the hepatic levels of IL-6, IL-1β and TNF-α, and then quantified the hepatic levels of AFP and CEA.

From these determinations, it could be seen that LPS injection induced a significant high expression of IL-6 (Fig. 1A), IL-1β (Fig. 1B) and TNF-α (Fig. 1C). Accordingly, the hepatic levels of AFP (Fig. 1D) and CEA (Fig. 1E) were significantly higher in LPS-injected mice than those in NC mice. These data indicated that the inflammatory response induced by LPS had a positive correlation with the expression of liver cancer markers.

**Figure 1.**
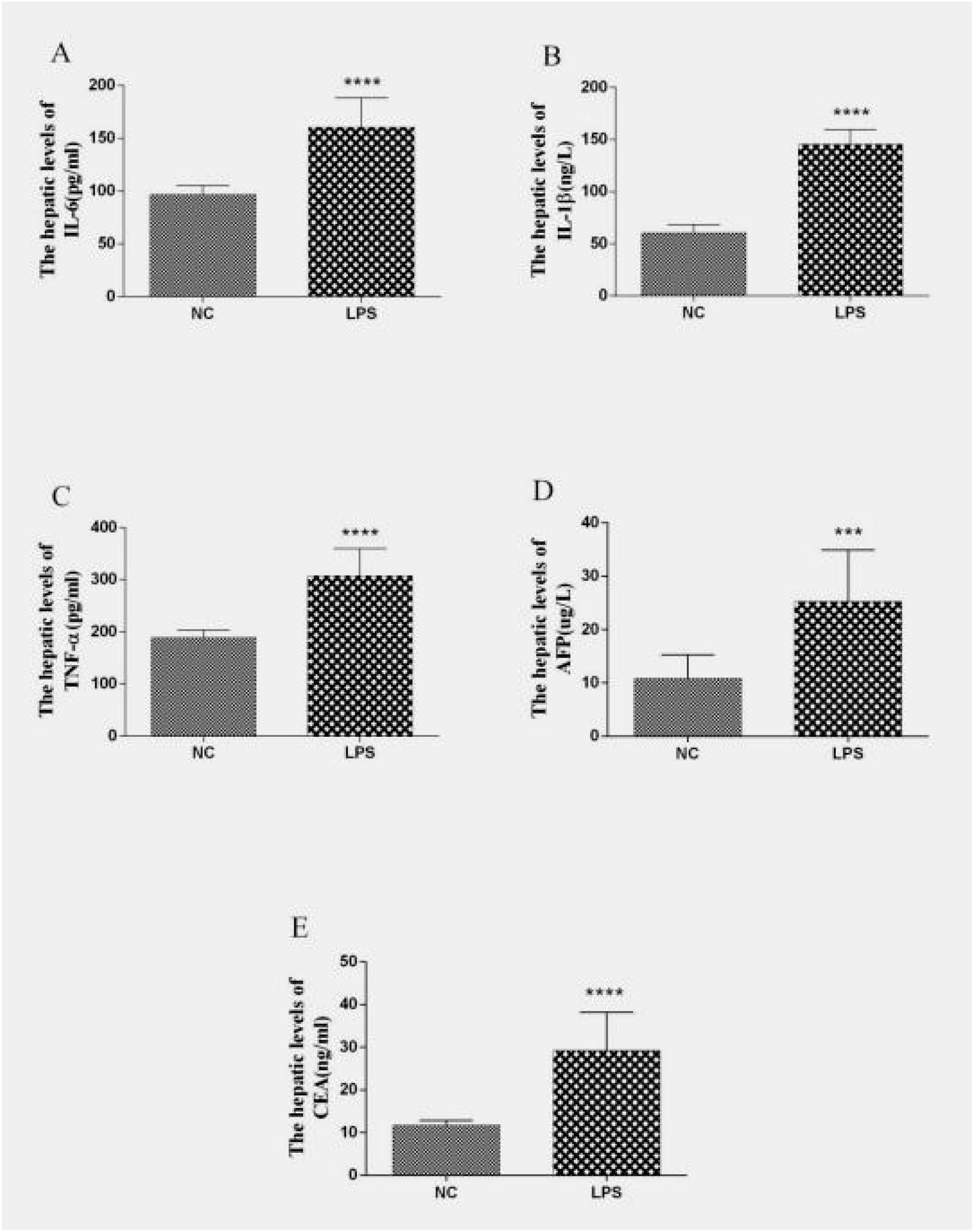
Effects of the inflammatory response on the expression of liver cancer markers in mice administered LPS. (A) The hepatic IL-6 level; (B) the hepatic IL-1β level; (C) the hepatic TNF-α level; (D) the liver cancer-specific tumor markers of AFP level; (E) the hepatic CEA level in NC and LPS mice were determined by ELISA. Results were normalized to scrambled. Data are presented as mean±standard deviation, all experiments are representative of three replicates. Compared with NC mice. *: *p* < 0.05, **: *p* < 0.01.

### BPDC reverses LPS-driven hepatic hyperplasia

To evaluate the histopathological changes in mice for 8-week LPS injections and for 3-week BPDC feeding, the fixed hepatic slices from each group were examined using HE staining. As shown in Fig. 2, compared with the sample of NC mice, LPS injection strikingly induced hepatic inflammatory infiltration and hyperplasia. Treatment of LPS-injected mice with BPDC remarkably attenuated LPS-driven hepatic inflammatory infiltration and hyperplasia compared with the sample of LPS mice. These results addressed that the long-term inflammation triggered by LPS could cause hepatic hyperplasia, and administration with BPDC could ameliorate LPS-induced hepatic hyperplasia.

**Figure 2.**
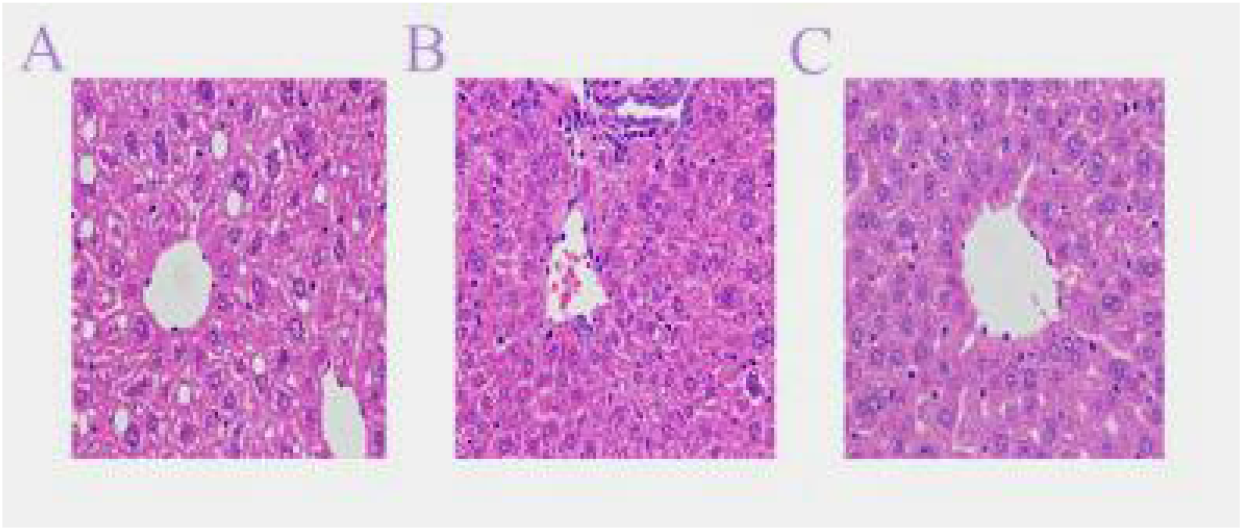
Effects of LPS-driven inflammatory responses on hepatic hyperplasia and BPDC-reversed histopathological changes in mice. The fixed hepatic samples were from NC, LPS, LPS+BPDC mice. (A) The hepatic sample from NC mice was examined by H&E (400×). (B) The hepatic sample from LPS mice was examined by H&E (400×). (C) The hepatic sample from LPS+BPDC mice was examined by H&E (400×).

### BPDC upregulates PTEN by decreasing miR-21

In order to evaluate the regulatory effect of BPDC on microRNAs, we measured the hepatic miR-21 levels in NC, LPS and LPS+BPDC mice. The results shown in Fig. 3A indicated that LPS injection increased hepatic miR-21 levels by approximately 3.87-fold compared with those of the hepatic sample of NC mice. Treatment with BPDC significantly attenuated the elevation of hepatic miR-21 levels compared with those of the hepatic sample of LPS mice.

**Figure 3.**
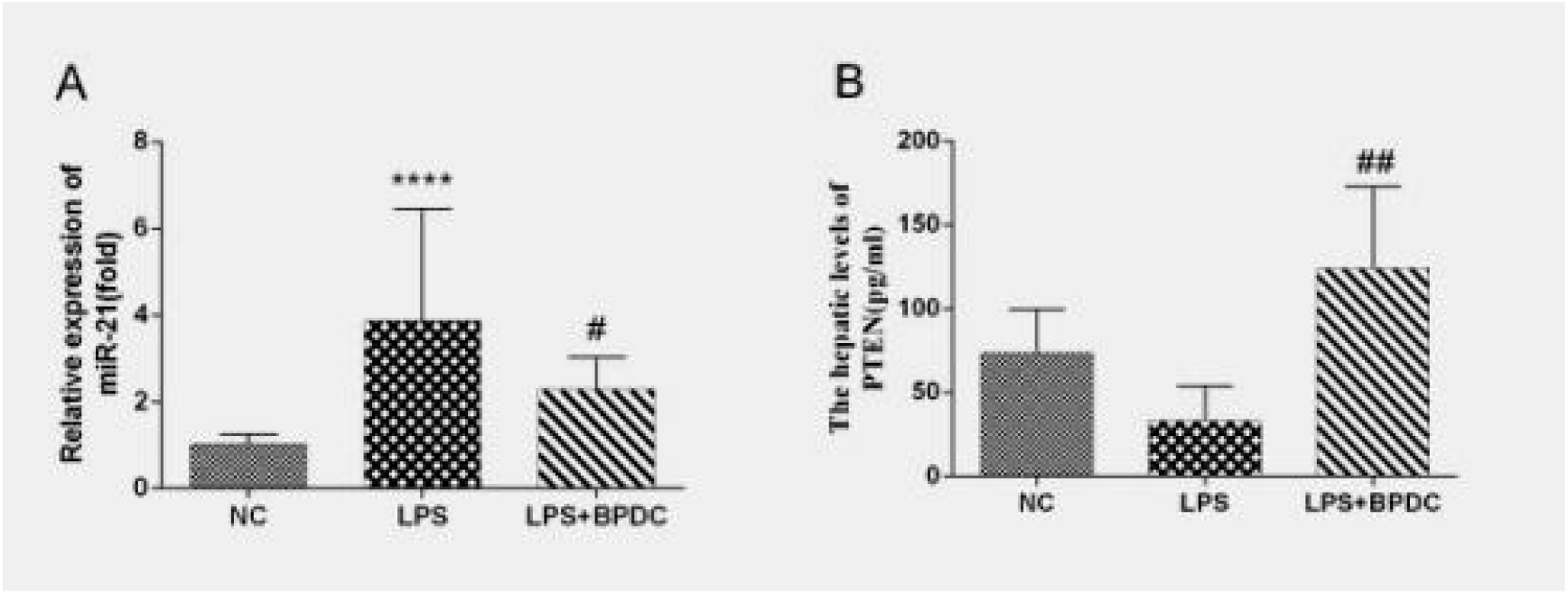
The regulatory effect of BPDC on LPS-driven miRNAs and target proteins in NC, LPS, LPS+BPDC mice. (**A**) miR-21 levels were measured by qRT-PCR; (**B**) PTEN levels were quantified by ELISA. Data are presented as mean ± standard deviation. *: *p* < 0.05, **: *p* < 0.01, compared with NC mice; #: *p* < 0.05, ##: *p* < 0.01, compared with LPS mice.

It is generally accepted that microRNAs do not encode the target proteins, but exert their function by regulating the expression of the downstream target genes. Thus, we identified the hepatic level of the potential target protein PTEN. These results indicated that a higher hepatic level of PTEN was measured in BPDC mice than in LPS mice (Fig. 3B).

In brief, these results unambiguously indicated that LPS-induced miR-21 levels a were predominantly elevated, whereas BPDC downregulated miR-21 and upregulated PTEN in LPS mice, suggesting a reverse correlation of miR-21 with PTEN.

### BPDC upregulates tumor suppressors via decreasing miR-122 and miR-24

To evaluate the effects of BPDC on LPS-driven miRNAs and target proteins, we determined the hepatic level of miR-122, miR-24, and the tumor suppressors, FOXO1 and MEN1, in NC, LPS, and LPS+BPDC mice. As consequences, LPS injection dramatically increased the hepatic levels of miR-122 (7.8-fold) and miR-24 (2.0-fold) compared to those in NC mice. Administration of BPDC reduced the hepatic levels of miR-122 and miR-24 compared with those of LPS mice (Fig. 4A and C). The expression levels of FOXO1 and MEN1 were lower in LPS mice compared with those of NC mice; administration of BPDC upregulated the hepatic levels of FOXO1 and MEN1 compared with those of LPS mice (Fig. 4B and D). These data indicated that LPS-induced miR-122 and miR-24 were expressed differentially, and BPDC exerted a therapeutic effect by upregulating FOXO1 and MEN1 after decreases in miR-122 and miR-24.

**Figure 4.**
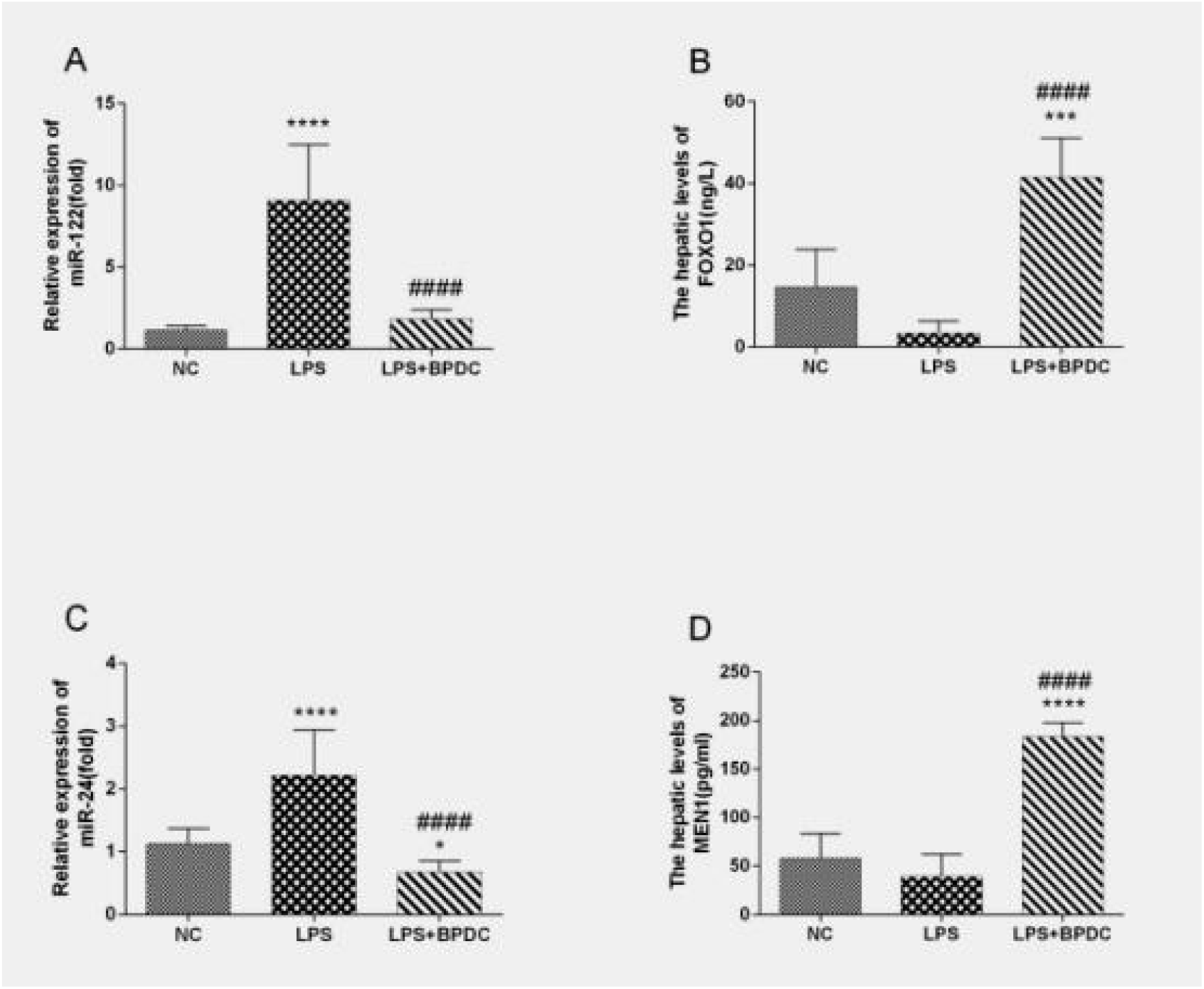
The effect of BPDC on LPS-driven miRNAs and target proteins in NC, LPS, LPS+BPDC mice. (A) miR-122 levels were measured by qRT-PCR. (B) FOXO1 levels were measured by ELISA. (C) miR-24 levels were quantified by qRT-PCR. (D) MEN1 levels were quantified by ELISA. Data are presented as mean ± standard deviation. *: *p* < 0.05, **: *p* < 0.01, compared with NC mice; #: *p* < 0.05, ##: *p* < 0.01, compared with LPS mice.

In order to figure out the regulation patterns of miRNAs on target proteins, we analyzed the correlation between the expression levels of miRNAs and target proteins in mice treated with LPS. As shown in Fig. 5, a significant inverse correlation of miR-21 levels with PTEN levels (r = 0.635, *P* = 0.0082), and miR-24 levels with MEN1 levels (r = 0.6439, *P* = 0.0096), but except for miR-122 levels and FOXO1 levels (r = 0.3786, *P* = 0.1819). These results indicated that there is a negative correlation between the selected miRNAs levels and corresponding target proteins levels. Further experiments warrant to elucidate the regulatory mechanism of miRNAs on its corresponding target proteins.

**Figure 5.**
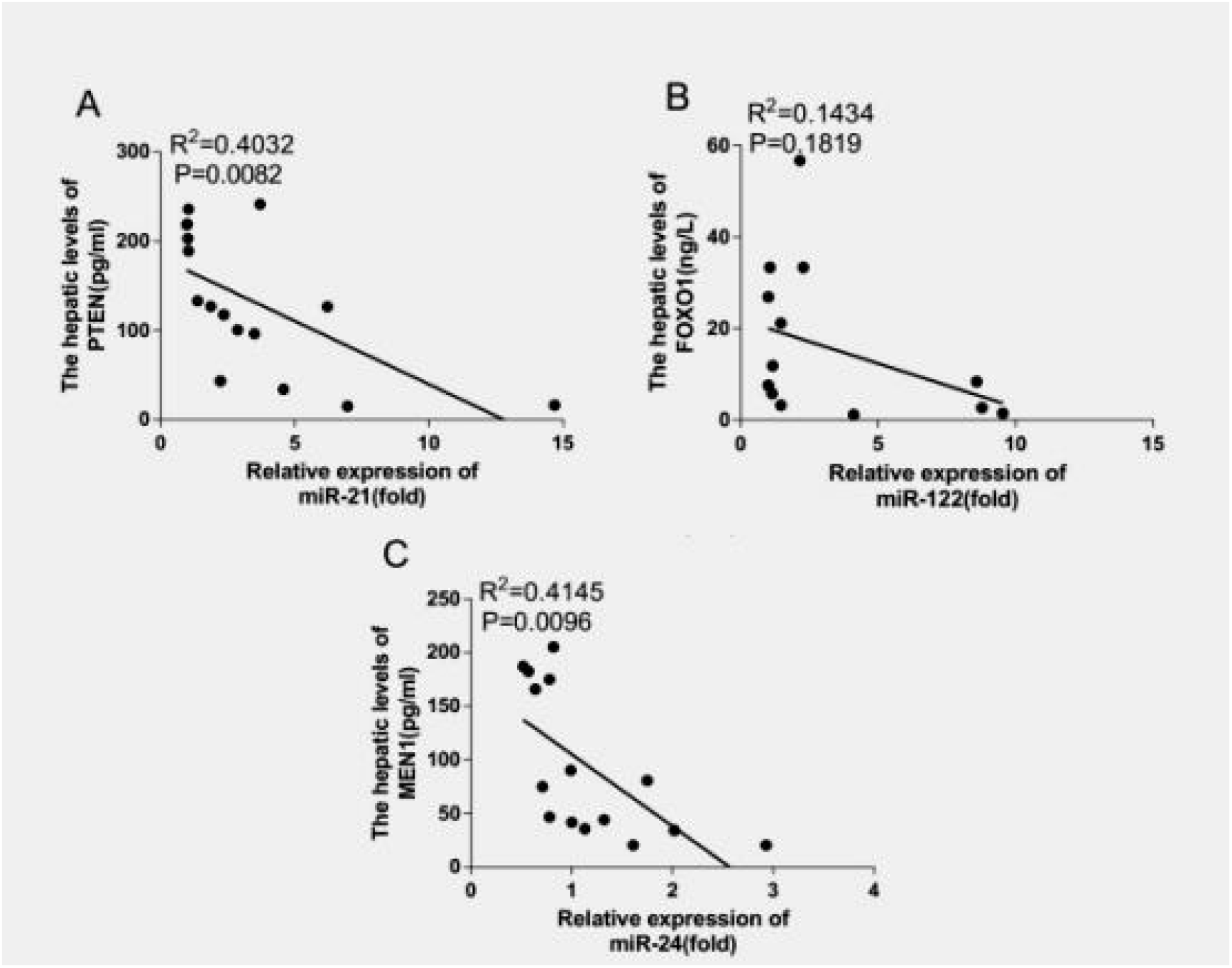
The correlation analysis of hepatic miRNAs and target proteins in mice given by LPS. (A) A correlation of the hepatic miR-21 levels with PTEN levels; (B) A correlation of the hepatic miR-122 levels with FOXO1 levels; (C) A correlation of the hepatic miR-24 with MEN1 levels analyzed using the software Graphpad prism 6.01. *p*<0.05 is statistically significant.

*BPDC fails to downregulate AEG-1 via increasing miR-195*. As shown above, BPDC exerted beneficial effects via upregulating tumor suppressors, PTEN, MEN1 and FOXO1. Whether does BPDC also down-regulate tumor markers? For this purpose, we quantified miR-195 and its downstream target protein, AEG-1,a tumor marker. We observed that the levels of miR-195 and AEG-1 in LPS mice were slightly higher than those in NC mice without significant difference. However, administration of BPDC failed to decrease AEG-1 levels compared with LPS mice (Figure 6). These results clearly indicated that BPDC exerted anti-tumor effects not by downregulating tumor markers.

**Figure 6.**
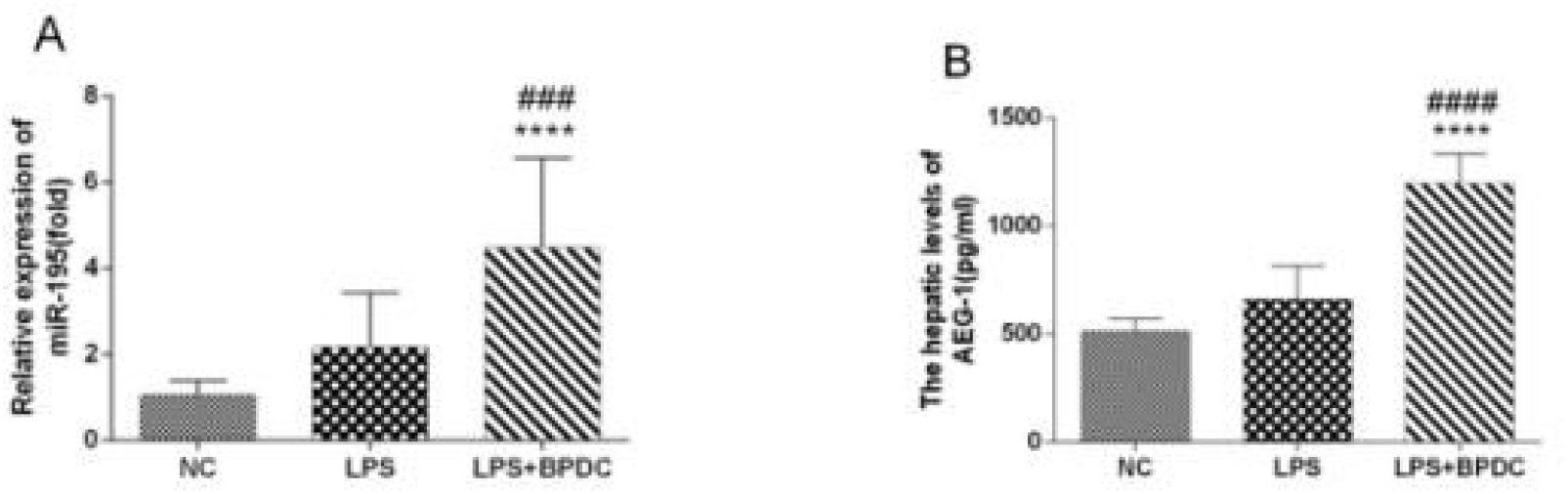
The correlation of BPDC-induced miRNAs with target protein levels in NC,LPS, LPS+BPDC mice. (A) miR-195 levels were measured by qRT-PCR. (B) AEG-1 levels were quantified.by ELISA. Data are presented as mean±standard deviation. *: *p* < 0.05, **: *p* < 0.01, compared with NC mice; #: *p* < 0.05, ##: *p* < 0.01, compared with the L mice.

## Discussion

A large number of epidemiological and clinical studies have shown a link between sustained inflammation and increased liver cancer risks, in which more than HCC occurs accounting for 90% in the context of liver injury and inflammation (25,26). In the present study, we injected mice with LPS for 8 weeks to mimic a chronic inflammatory milieu, resulting in significant upregulation of pro-inflammatory cytokines and overexpression of HCC-specific biomarkers. From these results, it was anticipated that LPS-driven chronic inflammation can promote HCC by repeatedly activating the immune response and downstream signaling. Indeed, inflammation triggers T cells and macrophages release ROS, leading to oxidative pathogen eradication and organ injuries (27,28). Oxidative stresses might result in DNA mutation, which is considered to be the primary underlying cause of cancer (29). These results highlight that chronic inflammation is closely correlated with tumorigenesis including HCC and other liver pathogenesis via oxidative stresses.

In recent years, mounting evidence has indicated that activation of oncogenes and loss of function or mutation of tumor suppressor genes are the putative molecular mechanism of tumor onset, and miRNAs expressed aberrantly are involved in physiological processes during tumor development (30,31). For example, miR-21 promotes cell invasion, migration, and growth via repression the expression of PTEN, MMP-2 and MMP-9 (32). MiR-122 promotes cell apoptosis and inhibits apoptosis by targeting FOXO and inducing Bcl-w (33,34). Upregulation of miR-24 promotes cell proliferation by targeting MEN1 and regulates cell growth and apoptosis by targeting BCL2 (35,36). MiR-195, a tumor suppressor, inhibits cell proliferation via targeting AEG-1 (37). We found in this study that LPS injection leads to a significant elevation of miR-21, miR-122 and miR-24, but a decline of the tumor suppressors, PTEN, FOXO1, and MEN1. However, LPS injection only allows insignificant elevation of miR-195 and the tumor marker AEG-1, indicating that LPS induces PTEN, FOXO1, and MEN1 rather than AEG-1. It would be meaningful to explore a therapeutic strategy targeting the above miRNAs and corresponding target proteins for liver cancer in further studies.

Furthermore, we confirmed an inverse correlation of miR-21 levels with PTEN levels, miR-24 levels with MEN1 levels, and miR-122 levels and FOXO1 levels with significance or insignificance. MiRNA can post-transcriptionally regulate gene expression by base pairing with the 3’-untranslated regions (3UTRs), or, less commonly, with the 5’-untranslated regions (5UTRs), of the target messenger RNAs (mRNAs) (38), resulting in translational repression or target cleavage (39). A specific miRNA can regulate multiple genes, while one gene may be regulated by a variety of miRNAs (40,41). The regulatory link can be established between miRNAs and corresponding target proteins in a network manner, which affects cell growth, proliferation, differentiation, apoptosis and tumor metastasis (42–44).

Since the abnormal expression of miRNAs has been involved in the regulation of tumor development, whether inhibition of upregulated miRNAs or recovery of downregulated miRNAs might reduce or even prevent tumor occurrence has sparked a challenging discussion. At present, several miRNA inhibitory agents, such as anti-sense oligonucleotides, locked nucleic acid (LNA) anti-miRNAs and miRNA sponges, have been widely investigated in miRNA-based target therapies. The anti-sense oligonucleotide, a single-stranded RNA molecule complementary to the target miRNA, exerts a role of competitive inhibition by preventing miRNA from interacting with the target gene. For instance, intravenous injection with antagomirs against miR-122 suppresses the endogenous miR-122, which elevates the expression level of a target gene (45). The LNA anti-miRNAs, modified by an extra methylene bridge connecting the 2’-O atom and the 4’-C atom ‘locks’ the ribose ring in a C3’-endo or C2’-endo conformation, exhibit the higher thermal stability and increases the hybridization properties of oligonucleotides (46,47). Based on LNA, miravirsen treatment for hepatitis C, targeting miR-122, was underway in the Phase II clinical trial (48). MiRNA sponge is a mRNA whose 3’ untranslated region (UTR) contains several miRNA target sites that stably bind to (RISC), keeping it away from natural mRNA targets and leading to a long-term inhibition of miRNA genes (49). Particularly, miRNA mimics have been widely used in the repair of tumor suppressor miRNAs to enhance the function of endogenous miRNAs and strengthen negative regulatory effects on the downstream target genes. For example, miRNA mimics for miR-495 can inhibit tumor proliferation in non-small cell lung cancer (NSCLC) (50).

In the present study, we reported for the first time that BPDC, a chemical compound other than a sequence fragment, upregulated tumor suppressors, PTEN, FOXO1 and MEN1 through decreasing tumorigenic miRNAs, miR-21, miR-122, and miR-24, functioning as a miRNA inhibitory agent. We also found that BPDC elevated miR-195, but failed to downregulate AEG-1, suggesting that BPDC exerted therapeutic effects as a enhancer of tumor suppressors, but not as a inhibitor of tumor markers or oncogenes. Our results demonstrated that long-term inflammation had a positive promoting effect on the initiation of HCC, and LPS injection strikingly induced inflammatory infiltration and hepatic hyperplasia via elevation the hepatic levels of miR-21, miR-122, and miR-24. In addition, BPDC exerted therapeutic effects as a miRNA inhibitory agent to upregulate the hepatic levels of PTEN, FOXO1 and MEN1 through decreasing the hepatic levels of miR-21, miR-122, and miR-24. These data suggested an important role of miR-21, miR-122, and miR-24 in the molecular etiology of HCC, and also provided a theoretical basis for potential molecular target-based early diagnosis and treatment of HCC. Meanwhile, This study had preliminarily elucidated the therapeutic mechanism of BPDC in liver hyperplasic diseases.

## Acknowledgments

We thank Mr. Chao Zhou in Haoma Biotechnology Co, Guangzhou, China for his technical assistance in H & E analysis.

## Funding

This work was supported by National Natural Science Foundation of China (81673861).

## Availability of data and materials

The datasets used and/or analyzed during the current study are available from the corresponding author on reasonable request.

## Authors’ contributions

- Li-Li Tan performed the experiments, analyzed the data, contributed reagents/materials/analysis tools, prepared figures and/or tables, reviewed drafts of the paper.
- Dong-Mei Chen performed the experiments.
- Qing-Ping Zeng conceived and designed the experiments, analyzed the data, wrote the paper, prepared figures and/or tables, reviewed drafts of the paper.
- Chang-Qing Li reviewed drafts of the paper.

All authors read and approved the final manuscript.

## Ethics approval and consent to participate

Animal procedures were in accordance with the animal care committee at the Guangzhou University of Chinese Medicine, Guangzhou, China (Certificate No. 44005800007512). The protocol was approved by the Animal Care Welfare Committee of Guangzhou University of Chinese Medicine.

## Patient consent for publication

Not applicable.

## Conflicts of interest

The authors declare that they have no conflict of interest, financial or otherwise.

